# A Dynamic Search Mechanism Enables APE1 to Identify AP-Sites in DNA

**DOI:** 10.64898/2026.01.06.697944

**Authors:** Kaitlin M. DeHart, Peyton N. Oden, Tyler M. Weaver, Mattew A. Schaich, Bennett Van Houten, Bret D. Freudenthal

## Abstract

Apurinic/apyrimidinic (AP) sites are among the most frequent DNA lesions, arising thousands of times per cell each day. These AP-sites threaten genomic stability and, if left unrepaired, can lead to mutagenesis and human disease. The essential base excision repair enzyme Apurinic/Apyrimidinic Endonuclease I (APE1) initiates repair by cleaving DNA at AP-sites, yet how APE1 efficiently locates and recognizes these lesions within vast excesses of undamaged DNA has remained poorly understood. Using single-molecule imaging, we show that APE1 employs a dynamic search strategy that integrates 1D and 3D diffusion to rapidly scan DNA. On non-damaged DNA, APE1 undergoes fast diffusion, enabling efficient interrogation of large genomic regions within a single binding event. Upon encountering an AP-site, APE1 transitions from a mobile search state into a stationary, lesion-bound complex that is retained at the site of damage. Additional experiments with APE1 variants demonstrate that the intrinsically disordered N-terminal domain of APE1 supports 1D diffusion, whereas residue R177 stabilizes APE1 at the AP-site once recognized, and the catalytic residues D210 and E96 facilitate enzyme release after cleavage. Together, these findings define the molecular basis by which APE1 balances rapid genome surveillance with stable lesion engagement and timely release. More broadly, this work provides a mechanistic framework for how DNA repair enzymes efficiently locate and process rare lesions embedded within an excess of undamaged DNA.

**Significance Statement:** DNA repair enzymes face the formidable challenge of locating rare lesions hidden within vast stretches of undamaged DNA. The essential enzyme APE1 is tasked with identifying and initiating repair of AP-sites, a frequent DNA lesion, to maintain genome integrity. We show that APE1 overcomes this challenge by using a dynamic search mechanism that integrates rapid scanning with precise lesion recognition. Its unstructured N-terminal domain facilitates fast movement along non-damaged DNA, while specific active site residues stabilize engagement at AP-sites and release after catalysis. This coordination of diffusion, recognition, and turnover allows APE1 to efficiently survey the genome and repair AP-sites to maintain genome stability.

## Introduction

DNA damage from endogenous metabolic processes and exogenous environmental factors presents a constant threat to genomic stability (1–3). Among the most prevalent DNA lesions are Apurinic/Apyrimidinic (AP) sites, which can form spontaneously up to 10,000 times per cell per day (4, 5). If unrepaired, AP sites can be propagated as mutations during replication or converted into more dangerous forms of DNA damage like single strand breaks, both of which are significant contributors to carcinogenesis (2–7). Thus, efficient and timely repair of AP-sites is critical for maintaining genome integrity and preventing disease (8–10).

In mammalian cells, the repair of AP-sites is initiated by Apurinic/Apyrimidinic Endonuclease 1 (APE1), a key enzyme in the Base Excision Repair (BER) pathway (11–13). To initiate repair of AP-sites, APE1 binds and flips the AP-site extrahelically into its active site (14–16). APE1 then cleaves the DNA phosphodiester backbone 5’ of the AP-site via a metal-dependent hydrolysis reaction (14–16). While the catalytic mechanism of APE1 at AP-sites has been well characterized through structural and kinetic studies (14–20), how APE1 efficiently locates AP-sites remains poorly understood. This is particularly relevant since AP-sites, though common compared to other DNA lesions, represent a comparatively small fraction of the approximately 3 billion base pairs in the human genome (4, 5, 21). The identification of AP-sites by APE1 therefore presents a complex search problem requiring this enzyme to both locate and cleave AP-sites for efficient repair.

To directly observe this process, we employed correlated optical tweezers with fluorescence microscopy (CTFM) to track fluorescently labeled Halo-tagged APE1 interacting with multiple DNA substrates at single-molecule resolution. We found that APE1 is diffusive on non-damaged DNA, suggesting it can efficiently scan large genomic regions. Introduction of a site-specific AP-lesion allowed us to distinguish between one-dimensional (1D) and three-dimensional (3D) search modes and to quantify APE1’s retention at AP-sites. To further dissect the underlying mechanisms, we generated three APE1-Halo mutants: APE1^R177A^-Halo, APE1^Dead^-Halo, and APE1^Δ1-42^-Halo. Mutation of R177, revealed that this residue is critical for stabilizing APE1 at AP-sites following recognition. In contrast, catalytic residues D210 and E96, mutated in our catalytically inactive construct, facilitate APE1 dissociation after cleavage. Finally, deletion of the N-terminal unstructured redox domain of APE1 (Δ1-42) revealed that this region is indispensable for 1D diffusion along non-damaged DNA. Collectively, these results support a model in which APE1 uses both 1D and 3D diffusion to scan the genome, with 1D search driven by the flexible N-terminal domain. Upon encountering an AP-site, APE1 engages the lesion and is retained via R177, while D210 and E96 promote release. Altogether, our findings and proposed model explain how APE1 efficiently locates AP-sites to drive an essential DNA repair process.

## Results

To investigate the damage search dynamics of APE1 for AP-sites, we purified recombinant APE1-Halo, confirmed its enzymatic activity, and fluorescently labeled the recombinant protein with JaneliaFluor Halo-tag ligand 552 (22) for use in a LUMICKS C-Trap CTFM system. We prepared a 12.65 kbp biotinylated DNA substrate consisting of either non-damaged DNA or DNA containing a single, central non-hydrolyzable AP-site analog (Fig. 1*A*). Streptavidin-coated polystyrene beads were captured in dual optical traps, allowing manipulation of individual DNA strands for visualization of APE1-Halo interactions (Fig. 1*B*). The DNA was visualized in 2D images known as kymographs in which a line across the position of the DNA is imaged at a frame rate of 15 ms. From our kymographs, we tracked the position of individual APE1-Halo molecules bound to DNA to determine their diffusivity, dwell times, and when applicable gap times (Fig. 1*C*).

**Figure 1:**
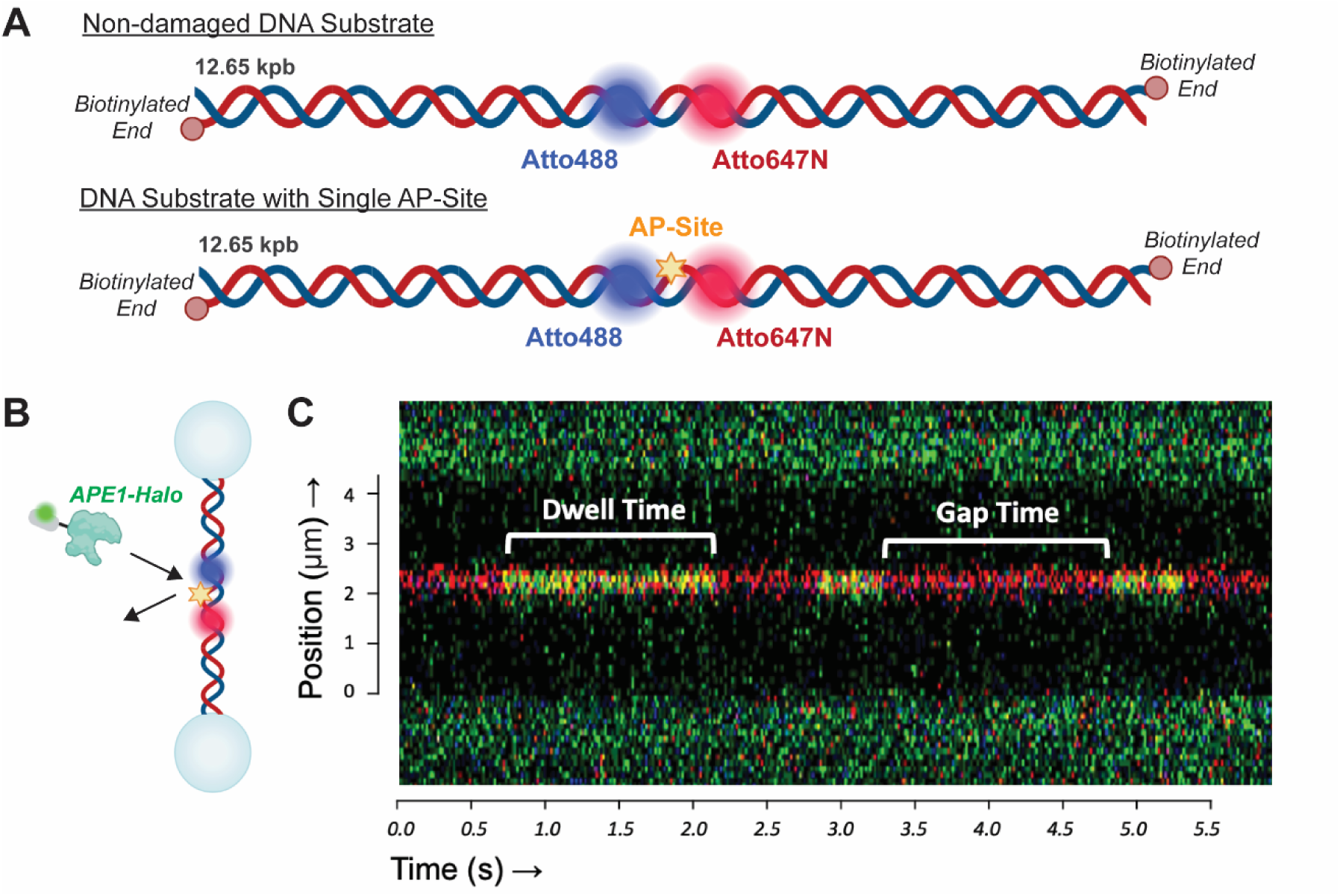
DNA Substrates and Their Visualization During CTFM Experiments. (A) The non-damaged and damaged DNA substrates used in CTFM experiments. Both substrates are identical except for a single, site specific THF AP-site analog on the damaged substrate. (B) Biotinylated DNA substrates are immobilized via streptavidin coated polystyrene beads captured with optical traps allowing binding interactions with protein in the surrounding solution. (C) An example kymograph demonstrating APE1 binding events. Examples of dwell times and gap times relative to binding are labeled. Parts of this figure were created with BioRender.com.

### APE1 Diffuses Along Non-Damaged DNA Through Rapid One-Dimensional Movement

To understand how APE1 searches non-damaged genomic DNA, we first observed APE1-Halo interacting with our non-damaged DNA substrate. Analysis of over 350 binding events revealed that the vast majority of APE1 molecules that interacted with non-damaged DNA were motile after binding the DNA. 84.1% of all APE1-Halo binding events to non-damaged DNA were motile, while only 2.2% of exhibited stationary behavior, and 13.6% exhibited a combination of both stationary and motile behavior (Fig. 2*A*). Given the largely motile behavior of APE1-Halo on non-damaged DNA, we hypothesized these motile events indicate search of non-damaged DNA for AP-sites, and we further investigated the events exhibiting motile behavior to determine the dynamics of a potential damage search mechanism.

**Figure 2:**
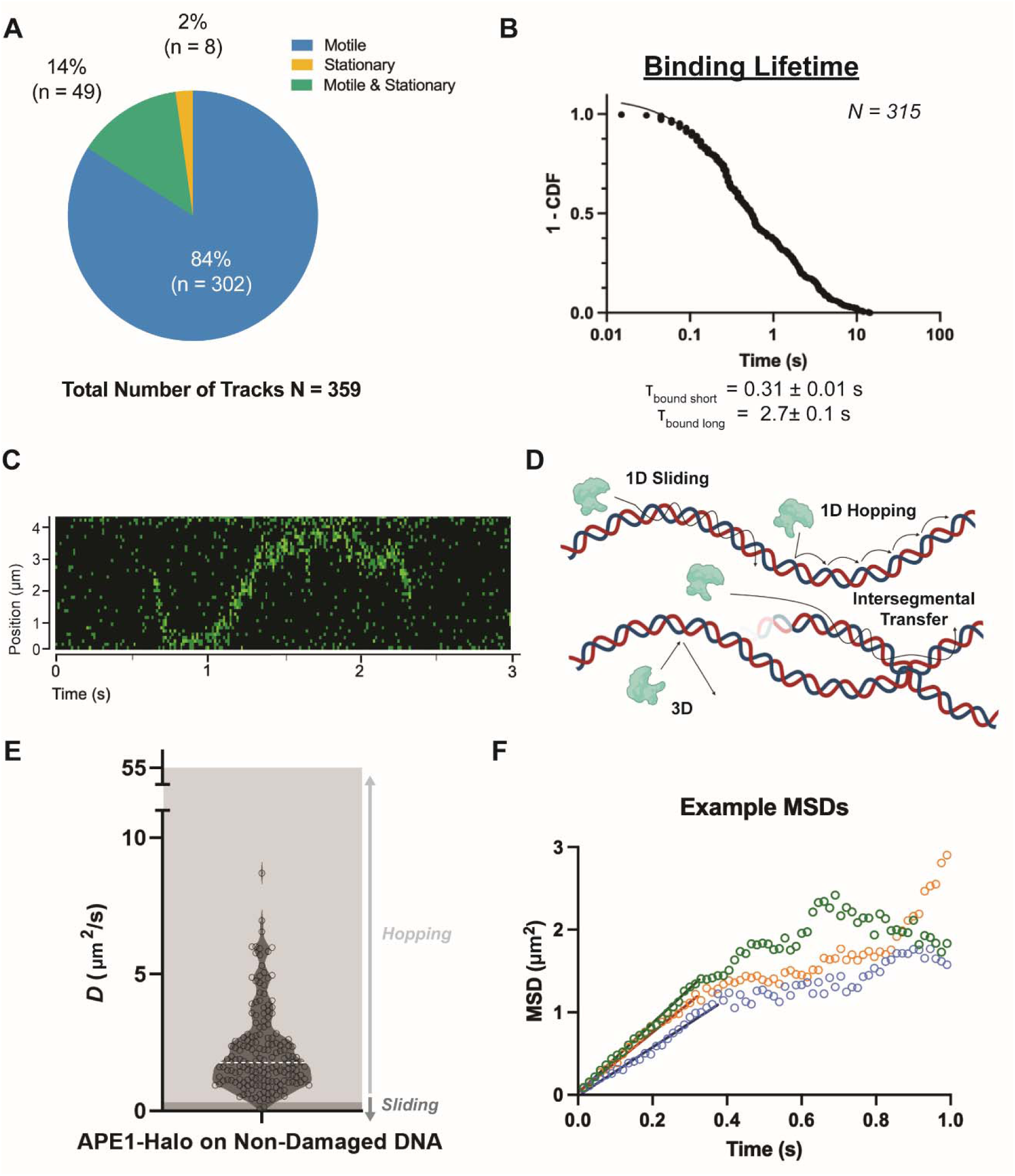
APE1-Halo is Motile and Diffusive on Non-Damaged DNA. (A) Movement behavior of APE1-Halo on non-damaged DNA. (B) 1-CDF plot of dwell times to determine binding lifetimes of short- and long-lived populations of APE1-Halo disassociating from non-damaged DNA. (C) An example kymograph showing APE1-Halo moving along non-damaged DNA. (D) Types of diffusion of proteins along DNA. (E) Violin plot of diffusion coefficients determined from MSD analysis of APE1-Halo on non-damaged DNA. The diffusion coefficients below the limit of sliding are indicated by a dark grey background, diffusion coefficients above the limit for sliding and below the limit for hopping are indicated by a light grey background, and the median diffusion coefficient is shown as a dashed white line. (F) Examples of MSD plots demonstrating the linear portion of MSD vs. time that is fit to determine the diffusion coefficient of a single binding event. Parts of this figure were created with BioRender.com.

We quantified the length of individual APE1-Halo binding events to non-damaged DNA (dwell times) and from these dwell times determined a representative binding lifetime (τ_bound_) using Poisson regression (Fig. 2*B*, Table 1) (23–25). This analysis revealed two types of motile APE1 populations, a shorter-lived population and a longer-lived population of APE1 on non-damaged DNA. The short population comprised 55% of the binding lifetime events, while the long population comprised 45% of events. The lifetime of the short population was 0.31 ± 0.01 s and the lifetime of the long population was 2.7 ± 0.1 s, respectively. Kinetic off-rate constants (*k_off_*), which are directly inversely proportional to binding lifetimes, are presented in *SI Appendix*, Table S1. Of note, we observed multiple individual APE1-Halo tracks capable of traversing the entirety of our 12.65 kbp non-damage DNA substrate within the time they were bound, suggesting APE1 is highly diffusive and can search significant amounts of DNA within a single binding event (Fig. 2*C*).

**Table 1:** Table of Binding Lifetimes and Gap Intervals of APE1-Halo and APE1-Halo Mutants. Summary of binding lifetimes (τ_bound_) and gap intervals (τ_free_) of APE1-Halo and mutants. Times are calculated for motile binding events on non-damaged DNA or for stationary binding events and the stationary portion of events exhibiting a mix of motile and stationary behavior at the AP-site. Times are separated into fast or slow populations, and the proportion of each population is given by distribution of the total population of binding events (N).

|  | <u>APE1-Halo</u> | <u>APE1<sup>R177A</sup>-Halo</u> | <u>APE1<sup>Dead</sup>-Halo</u> | <u>APE1<sup>Δ1-42</sup>-Halo</u> |
| --- | --- | --- | --- | --- |
| **Motile Binding Events on Non-Damaged DNA** |  |  |  |  |
| *Binding Lifetimes* |  |  |  |  |
| $\tau_{\text{bound short}}$ (s) | 0.31 ± 0.01 | | 0.47 ± 0.04 | |
| $\tau_{\text{bound long}}$ (s) | 2.7 ± 0.1 | | 1.2 ± 1.4 | |
| N | 315 |  | 334 |  |
| Distribution<sub>bound</sub> | 55.4% Short;<br>44.6% Long |  | 88.5% Short;<br>11.5% Long |  |
| **Stationary Binding Events at the AP-Site** |  |  |  |  |
| *Binding Lifetimes* |  |  |  |  |
| $\tau_{\text{bound short}}$ (s) | 0.75 ± 0.04 | 0.10 ± 0.01 | 0.44 ± 0.02 | 0.71 ± 0.03 |
| $\tau_{\text{bound long}}$ (s) | 5.7 ± 0.1 | 0.95 ± 0.05 | 22 ± 1 | 1.6 ± 0.1 |
| N | 381 | 406 | 266 | 1158 |
| Distribution<sub>bound</sub> | 17.6% Short;<br>82.4% Long | 80.4% Short;<br>19.6% Long | 26.9% Short;<br>73.1% Long | 47.0% Short;<br>53% Long |
| *Gap Intervals* |  |  |  |  |
| $\tau_{\text{free short}}$ (s) | 3.0 ± 0.1 | 3.9 ± 0.1 | 8.7 ± 0.3 | 2.0 ± 0.1 |
| $\tau_{\text{free long}}$ (s) | 24 ± 1 | 14 ± 1 | 71 ± 7 | 4.7 ± 0.8 |
| N | 328 | 109 | 366 | 1086 |
| Distribution<sub>free</sub> | 45.2% Short;<br>54.8% Long | 60.1% Short;<br>39.9% Long | 57.9% Short;<br>42.1% Long | 83.0% Short;<br>17.0% Long |

We next quantified the diffusivity of APE1-Halo on non-damaged DNA to investigate whether these motile events exhibited a rapid search for damage. DNA binding proteins exhibit multiple modes of diffusivity on DNA that include one-dimensional (1D) diffusivity, three-dimensional diffusivity (3D), and intersegmental transfer (Fig. 2*D*) (26–30). While we are unable to observe intersegmental transfer in our system, we did observe both 1D and 3D diffusivity. Here we define molecules of APE1-Halo that bound to non-damaged DNA and were motile as having 1D diffusion, while molecules that bound DNA and remained stationary are defined as having 3D diffusion. We did not appreciably observe 3D diffusive events on non-damaged DNA. Comparatively, APE1-Halo readily exhibited 1D diffusion on non-damaged DNA and we therefore sought to quantify APE1-Halo’s 1D diffusivity. To do so, we calculated the mean squared displacement (MSD) of individual binding events to obtain individual diffusion coefficients (*D*) and then determined an median diffusion coefficient (*D_med_*) from the full population of individual diffusion coefficients (Fig. 2*E-F*). The median diffusion coefficient of APE1-Halo on non-damaged DNA was 1.8 ± 1.9 µm^2^/s. To our knowledge and using this approach, this is one of the fastest measures of 1D diffusivity observed for a DNA-binding protein moving along DNA (24, 25, 31, 32). Furthermore, taking this median diffusion coefficient of 1.8 ± 1.9 µm^2^/s and a weighted average of both populations of binding lifetimes of 1.4 ± 0.1 s, APE1 would be capable of theoretically scanning 6.4 ± 3.5 kbp of DNA in a single binding event.

1D diffusivity can be further broken into two modes of binding, either 1D movement that is rotationally coupled to the structure of the DNA helix also known as “sliding” or non-rotationally coupled 1D movement known as “hopping” (Fig. 2*D*) (25, 28–30). To determine whether the mode of APE1-Halo’s 1D diffusion was restricted to either sliding or hopping, we next calculated the theoretical maximum diffusion coefficients (*D_max_*) of APE1-Halo if it were either sliding or hopping on non-damaged DNA. While sliding and hopping are both types of 1D diffusion, the friction or “drag” of a protein along DNA is greater when 1D movement is rotationally coupled (Eq. 5) than when it is non-rotationally coupled (Eq. 4) (28). We estimated the maximum diffusion coefficient of APE1-Halo when sliding as 0.29 µm^2^/s, and the maximum diffusion coefficient of APE1-Halo when hopping as 55 µm^2^/s. Our median diffusion coefficient for APE1-Halo is above the limit for sliding and within the limit for hopping. The majority of our individual diffusion coefficients are also above the limit for sliding, though some events are still below the theoretical limit for sliding. Overall, our results indicate that APE1-Halo does both sliding and hopping on non-damaged DNA. From these results, we propose that APE1 rapidly diffuses along non-damaged DNA in a 1D manner using hopping and/or sliding to search for damage.

### APE1 Recognizes and Stably Engages AP-Sites During the Search Process

Having defined the behavior of APE1 on non-damaged DNA, we next examined how APE1 behaves upon encountering an AP-site. To do so, we monitored APE1-Halo interactions with DNA containing a single, centrally positioned non-hydrolyzable tetrahydrofuran (THF) AP-site analog (Fig. 3*A*). In contrast to its behavior on non-damaged DNA, APE1-Halo exhibited a marked increase in stationary binding events on AP-DNA, accompanied by a reduction in exclusively motile behavior. Specifically, 39.6% of binding events were stationary, 38.8% were exclusively motile, and 21.6% displayed a combination of motile and stationary behavior (Fig. 3*B*). Notably, for events exhibiting stationary behavior, over 90% of these events localized to the AP-site, indicating specific lesion recognition and stable engagement by APE1.

**Figure 3:**
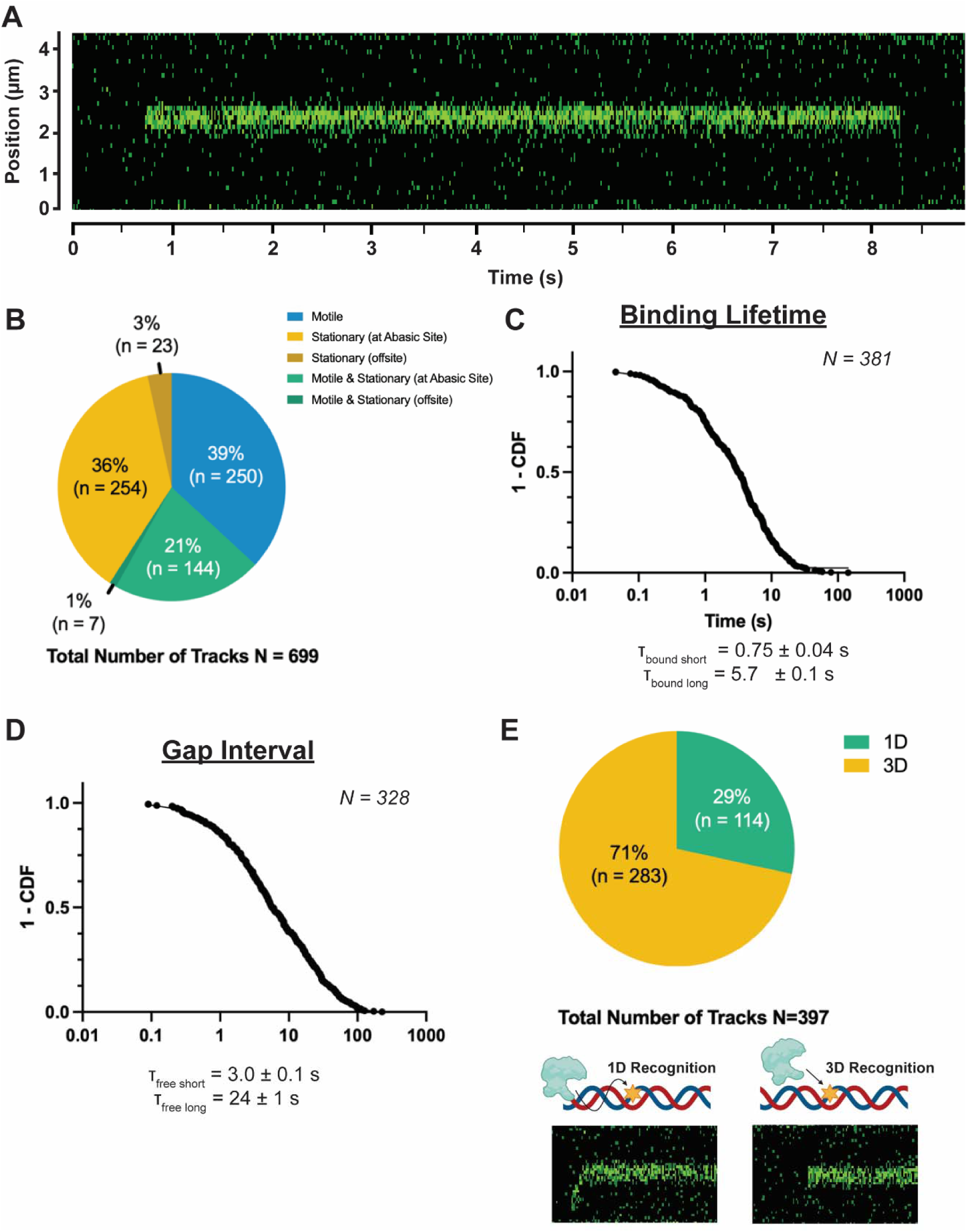
APE1-Halo Recognizes and is Retained at AP-Sites. (A) Exemplary kymograph of APE1-Halo recognizing and being retained at the centrally located AP-site. (B) Movement behavior of APE1-Halo on AP DNA. (C) 1-CDF plot of dwell times to determine the binding lifetimes of short- and long-lived populations of APE1-Halo disassociating from AP DNA. (D) 1-CDF plot of gap times to determine gap intervals of APE1-Halo binding to AP DNA. (E) Recognition of the AP-site by APE1-Halo via 1D diffusion or 3D diffusion. Parts of this figure were created with BioRender.com.

The events at the AP-site persisted substantially longer than motile events on non-damaged DNA, consistent with stable lesion engagement by APE1. For stationary binding events, as well as the stationary portion of the mixed motile and stationary events at the AP-site, we observed short-lived and long-lived binding lifetime populations (Fig. 3*C*, Table 1). The short-lived population comprised only 18% of lifetimes, while the long-lived population comprised 82% of lifetimes. The binding lifetime of the short population was 0.75 ± 0.04 s, while the lifetime of the long population was 5.7 ± 0.1 s. This represents a substantially increased lifetime of APE1 on damaged DNA when compared to our non-damaged lifetime.

We also determined the gap interval from individual gap times between successive APE1 binding events at the AP-site to assess the binding dynamics of lesion engagement (Fig. 3*D*, Table 1). The gap interval (τ_free_) similarly revealed two populations, with a short population (45% of events) exhibiting a gap interval of 3.0 ± 0.1 s and a longer population (55% of events) with a gap interval of 24 ± 1 s. Since we were able to determine both binding lifetimes and gap intervals for APE1-Halo at the AP-site, we were also able to estimate a dissociation rate constant (*K*_D_) from the corresponding on and off rate constants. To account for both populations contributing to each rate constant, we calculated weighted average on and off rate constants reported in *SI Appendix*, Table S2. Using these weighted average rate constants , we estimated a binding affinity (*K*_D_) of 0.55 ± 0.04 nM for APE1-Halo at an AP-site (*SI Appendix*, Table S2), in close agreement with our ensemble EMSA measurements and previously reported published values (*SI Appendix*, Fig S1*B-C*) (17, 33). Collectively, these values demonstrate that APE1 transitions from a transient, highly mobile search state on non-damaged DNA to a long-lived, stably engaged complex upon encountering an AP-site.

To determine how APE1 identifies AP-sites during its search process, we analyzed whether lesion recognition occurred via 1D along DNA or through direct 3D binding from solution (Fig. 3*E*). We found that APE1-Halo recognized AP-sites predominantly through 3D diffusion, accounting for 71% of all binding events at the lesion, while the remaining 29% of events involved APE1 locating the AP-site through 1D diffusion along the DNA. These results indicate that although APE1 is capable of searching DNA through rapid 1D scanning, direct binding from solution represents a major route of lesion recognition under our experimental conditions. Importantly, regardless of whether AP-sites were identified through 1D or 3D diffusion, APE1 transitioned into a stationary, long-lived complex upon lesion engagement. This convergence on a common, stably bound state suggests that either search pathway ultimately funnels APE1 into the same productive lesion bound conformation required to facilitate repair.

### Active Site Residues Regulate APE1 Retention and Release at AP-Sites

To define the molecular features that regulate APE1 engagement and release at AP-sites following lesion recognition, we examined the contributions of key active site residues with distinct functional roles. Structural and kinetic studies have shown that residue R177 of APE1 forms a stabilizing clasp at the AP-site and mutations of this residue increase dissociation of APE1 after lesion engagement (14, 15, 34). In addition, the catalytic residues D210 and E96 coordinate active site chemistry and are essential for APE1 catalysis (14, 15, 35). We therefore used single-molecule imaging to directly test how mutation of R177 and mutation of D210 and E96 differentially alters APE1 mobility, lesion engagement, and retention at AP-sites during the search and recognition process.

We first characterized our APE1^R177A^-Halo mutant on DNA containing an AP-site (Fig. 4*A*). In contrast to wild-type APE1, APE1^R177A^-Halo binding events were predominantly stationary with 83% of events classified as stationary and only a small fraction exhibiting motile or mixed motile stationary behavior (Fig. 4*B*). Importantly, the majority of stationary events localized to the AP-site, with 84% occurring specifically at the lesion, indicating that mutation of R177 does not prevent AP-site recognition. However, despite this apparent AP-site engagement, APE1^R177A^-Halo was poorly retained at AP-sites compared to wild type APE1 (Fig. 4*C*, Table 1). Binding lifetime analysis revealed a dominant short population comprising 80% of events with an lifetime of 0.10 ± 0.01 s, while a minor (20%) longer population exhibited a lifetime of 0.95 ± 0.05 s. These lifetimes were substantially shorter than those observed for APE1-Halo, demonstrating that R177 is essential for stabilizing APE1 at AP-sites following lesion engagement.

**Figure 4:**
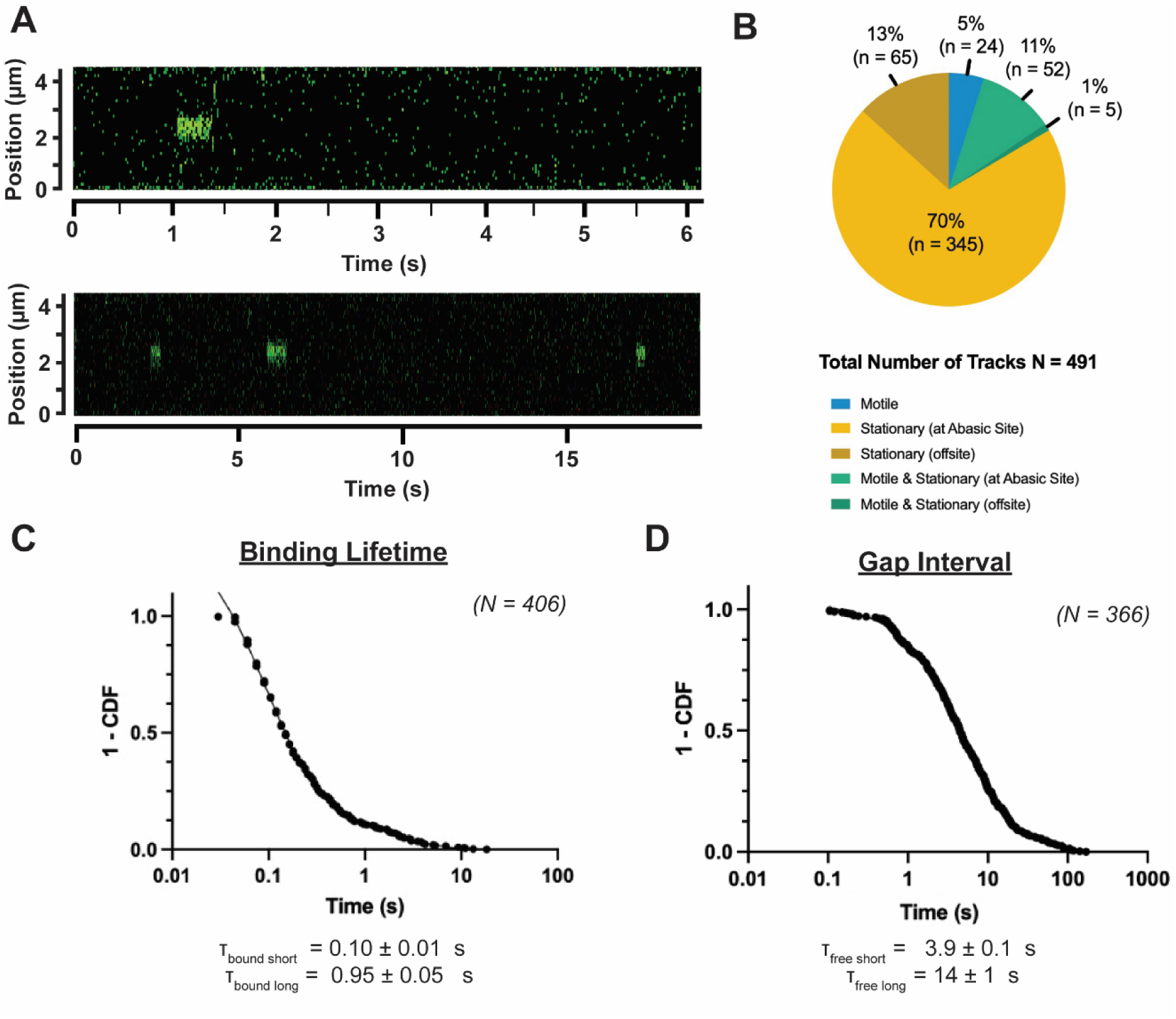
R177 Mediates APE1 Retention at AP-Sites. (A) Exemplary kymographs of APE1^R177A^-Halo on AP DNA. (B) Pie chart of APE1^R177A^-Halo movement behavior on AP DNA. (C) 1-CDF plot of dwell times to determine short- and long-lived populations of binding lifetimes of APE1^R177A^-Halo disassociating from AP DNA. (E) 1-CDF plot of gap times to determine gap intervals of APE1^R177A^-Halo binding to AP DNA.

We next examined the binding dynamics of APE1^R177A^-Halo at AP-sites by analyzing the gap interval (Fig. 4*D*, Table 1). Analysis revealed two kinetic populations of gap intervals, with a short population comprising 60% of events and a longer population comprising 40% of events. The short population exhibited a time of 3.9 ± 0.1 s, while the longer population exhibited a time of 14 ± 1 s. Consistent with these association dynamics and markedly accelerated dissociation, the apparent binding affinity of APE1^R177A^-Halo for an AP-site was estimated to be 11 ± 1 nM (SI Appendix, Table S2), representing an approximately order of magnitude reduction relative to APE1-Halo. Together, these results indicate that mutation of R177 does not substantially impair AP-site engagement but instead destabilizes the lesion-bound complex by accelerating dissociation.

We next observed our APE1^Dead^-Halo mutant (Fig. 5*A*), which contains mutations to active site residues D210, which deprotonates the nucleophilic water, and E96, which stabilizes the Mg^2+^ enzymatic co-factor (14, 15, 35). While these two residues have predominantly been of interest to prevent AP-site cleavage (35, 36), a recent study found this mutant is retained at AP sites in chromatin, prompting our interest in these residues possible role in mediating APE1 retention on AP DNA (37). In our CTFM experiments with APE1^Dead^-Halo in the presence of AP-DNA, we observed a mixture of motile, stationary, and mixed motile/stationary binding events similar to those for APE1-Halo (Fig. 5*B*). Notably, only 10% of events were stationary, whereas 77% were motile and 13% exhibited a mixed behavior. Despite this lower fraction of stationary events, once APE1^Dead^-Halo engaged at the AP-site it remained bound for substantially longer than APE1-Halo. We expect the proportion of stationary events to be lower, at least in part, because once APE1^Dead^-Halo engaged the AP-site it remained bound for extended durations, limiting re-engagement and therefore reducing the total number of discrete stationary events observed. Consistent with this interpretation, analysis of binding lifetimes revealed two populations, with a short population comprising 27% of events and exhibiting a lifetime of 0.44 ± 0.017 s, and a dominant long population comprising 73% of events with a lifetime of 22.2 ± 1 s, approximately four times longer than APE1-Halo (Fig. 5C, Table 1).

**Figure 5:**
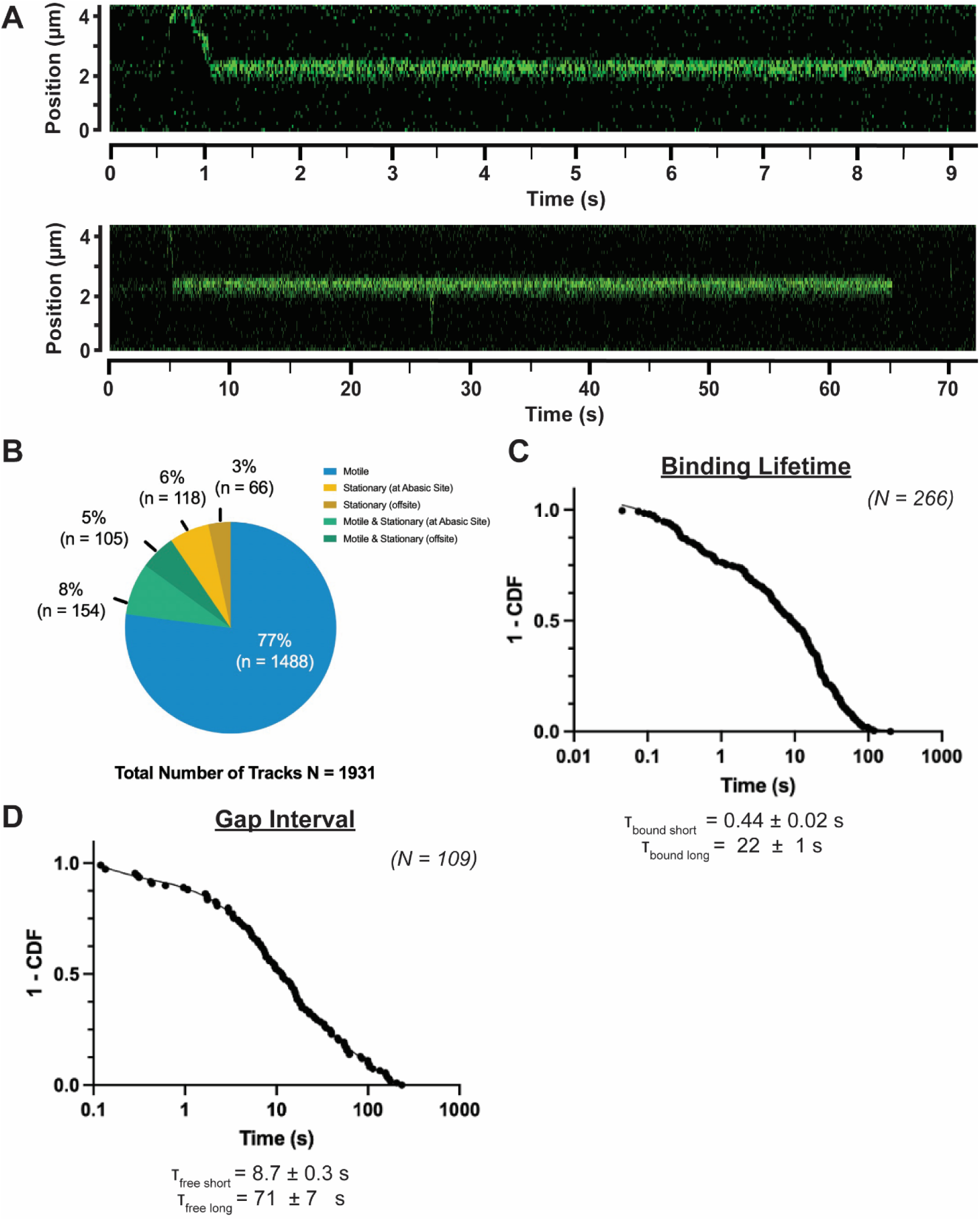
APE1^Dead^-Halo is Retained at AP-Sites Longer than APE1-Halo and Engages Less Often. (A) Exemplary kymographs of APE1Dead-Halo recognizing and being retained at the centrally located AP-site. (B) Movement behavior of APE1Dead-Halo on AP DNA. (C) 1-CDF plot of dwell times to determine binding lifetimes of short- and long-lived populations of APE1Dead-Halo disassociating from AP DNA. (D) 1-CDF plot of gap times to determine gap intervals APE1Dead-Halo binding to AP DNA.

We next examined the binding dynamics of APE1^Dead^-Halo at AP-sites by analyzing the gap interval (Fig. 5*D*, Table 1). Analysis of gap times revealed two gap interval populations, with a short population comprising 58% of events and a long population comprising 42% of events. The short population exhibited a gap interval of 8.7 ± 0.3 s, whereas the long population exhibited a markedly extended gap interval of 71 ± 7 s. Our single molecule kinetic binding characterization from the on and off rates of APE1^Dead^-Halo indicated that APE1^Dead^-Halo tightly binds AP-sites with an estimated affinity of 2.2 ± 0.2 nM (*SI Appendix*, Table S2) reflecting tight binding driven by the markedly reduced dissociation rate (*SI Appendix*, Table S1) and consistent with our ensemble binding measurements (*SI Appendix*, Fig. S1). Together with the prolonged binding lifetimes, these results demonstrate that mutation of D210 and E96 strongly stabilizes the lesion-bound complex while impairing productive release following lesion engagement. Notably, inhibiting APE1 cleavage is not sufficient to induce the binding lifetimes and gap intervals observed for our APE1^Dead^-Halo mutant since we also observed APE1-Halo on non-hydrolysable AP DNA.

### The APE1 N-Terminus Regulates 1D Diffusivity and AP-Site Recognition and Retention

Having defined how active site residues regulate APE1 stabilization and release at AP-sites, we next examined the contribution of the APE1 N-terminal domain to DNA binding and lesion search. The N-terminus of APE1 is intrinsically disordered and has been implicated in redox signaling and sequence-specific DNA interactions but is generally considered dispensable for AP-site cleavage (38–42). Whether this region contributes to the dynamic search process by which APE1 interrogates non-damaged DNA, however, has remained unclear. We therefore investigated the behavior of an APE1^Δ1-42^-Halo mutant using single-molecule imaging to determine how loss of the N-terminus affects APE1 diffusivity, lesion recognition, and engagement with AP-sites.

To determine whether the APE1 N-terminus contributes to binding and diffusion on non-damaged DNA, we examined our APE1^Δ1-42^-Halo mutant in our CTFM experiments. At our standard concentration of 250 pM APE1^Δ1-42^-Halo we did not detect any binding events to non-damaged DNA. Increasing the protein concentration ten-fold and monitoring potential interactions under flow similarly failed to reveal detectable binding events (Fig. 6*A*). To determine whether this reflected a reduced affinity rather than complete loss of binding, we next measured DNA binding by electrophoretic mobility shift assays (EMSAs). Whereas APE1-Halo bound non-damaged DNA with an apparent affinity of 54 ± 11 nM, deletion of the N-terminus reduced binding affinity to 83 ± 25 nM (SI Appendix, Fig. S1). Because loss of the APE1 N-terminus reduced the affinity of APE1 for non-damaged DNA, we expect it is important for APE1 binding to non-damaged DNA. Importantly, our data indicates that the APE1 N-terminus is critical in mediating the 1D search of APE1 as it diffuses along non-damaged DNA.

**Figure 6:**
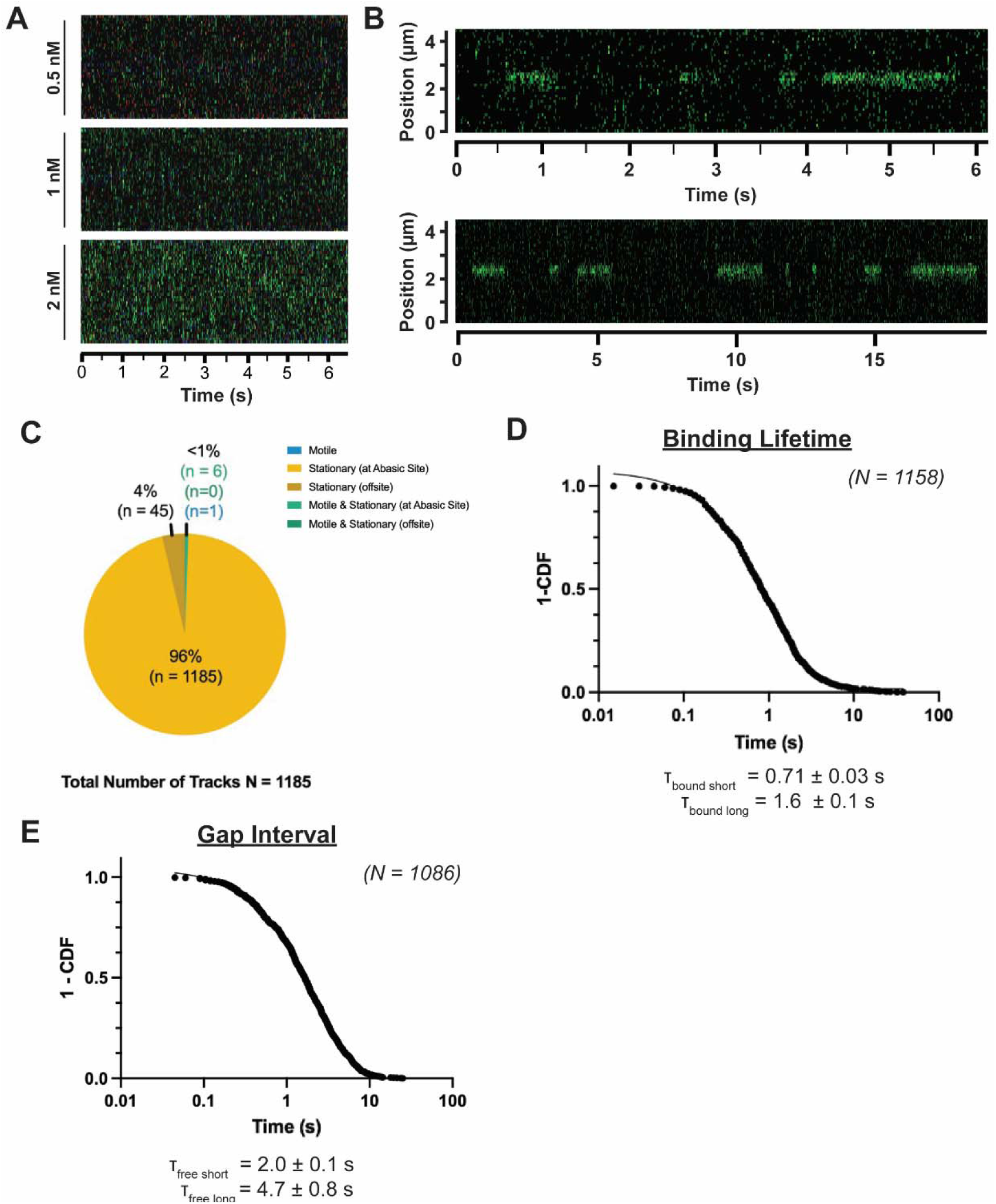
The N-Terminus Controls 1D Diffusion and AP-Site Engagement and Retention. (A) Exemplary kymographs of APE1Δ1-42A-Halo with non-damaged DNA. No specific binding events were observed even with increased APE1Δ1-42A-Halo concentration. (B) Exemplary kymographs of APE1Δ1-42A-Halo on AP DNA. (C) Pie chart of APE1Δ1-42A-Halo movement behavior on AP DNA. (D) 1-CDF plot of dwell times to determine binding lifetimes of short- and long-lived populations of APE1Δ1-42A-Halo disassociating from AP DNA. (E) 1-CDF plot of gap times to determine gap intervals of APE1Δ1-42A-Halo binding to AP DNA.

We next examined the behavior of APE1^Δ1-42^-Halo on DNA containing a single AP-site. In contrast to its inability to bind non-damaged DNA, APE1^Δ1-42^-Halo readily associated with AP-DNA in our CTFM experiments (Fig. 6*B*). Strikingly, nearly all observed binding events were stationary, with over 99% of events lacking detectable motile behavior and 96% of these stationary events localized specifically to the AP-site (Fig. 6*C*). Consistent with this behavior, APE1^Δ1-42^-Halo identified AP-sites almost exclusively through 3D diffusion, with fewer than 1% of events arising from 1D diffusion along DNA. These observations indicate that the APE1 N-terminus is dispensable for direct AP-site recognition but is required for 1D diffusion and lesion search along non-damaged DNA.

To quantify the binding dynamics of APE1^Δ1-42^-Halo at the AP-site, we analyzed dwell and gap times for stationary binding events (Fig. 6*D–E*, Table 1). Binding lifetime analysis revealed two populations, with a short population comprising 47% of events and exhibiting a lifetime of 0.71 ± 0.03 s, and a longer population comprising 53% of events with a lifetime of 1.6 ± 0.1 s. This effect was similar, but far more drastic than the one we observed for APE1^R177A^-Halo indicating that the N-terminus is also involved in APE1 retention despite no known direct interactions with the AP-site. Interestingly, gap interval analysis revealed rapid AP-site engagement, with a short population comprising 83% of events and exhibiting a gap interval of 2.0 ± 0.1 s, and a long population comprising 17% of events with a gap interval of 4.7 ± 0.8 s. Despite these changes in association and dissociation dynamics, the apparent binding affinity of APE1^Δ1-42^-Halo for an AP-site was estimated to be 0.65 ± 0.12 nM (*SI Appendix*, Table S2), similar to that of APE1-Halo. This suggests previous ensemble measurements may have missed the large differences in association and dissociation dynamics between these two mutants due to the similar binding affinity using ensemble approaches. Together, these findings demonstrate that the APE1 N-terminal domain is required for productive engagement with non-damaged DNA and one-dimensional diffusion during lesion search, while also modulating the kinetics of AP-site engagement and retention once a lesion is encountered.

## Discussion

The ability of DNA repair enzymes to search for and repair DNA lesions in a sea of undamaged DNA within the genome is a fundamental process safeguarding against the consequences of DNA damage. While the catalytic mechanisms of these enzymes during repair are generally well-studied (14–17), the mechanisms by which repair enzymes search the genome for sites in need of repair are less well understood. In this work, we demonstrate how the essential DNA repair enzyme APE1 searches for AP-sites and reveal the molecular mechanisms that this enzyme utilizes to facilitate its search for AP-site damage.

Our work here demonstrates an important role for APE1 binding interactions with non-damaged DNA during the DNA repair functions of APE1. These binding interactions with non-damaged DNA allow for 1D diffusion along non-damaged DNA to search for AP-sites. Observed 1D diffusion of APE1 along non-damaged DNA followed expected characteristics of a random walk (Brownian motion) (28). Furthermore, 1D diffusion of APE1 on non-damaged DNA could be broken down into a mix of both rotationally coupled “sliding” and non-rotationally coupled “hopping” 1D diffusion (25, 28–30). This mix of sliding and hopping behavior has been observed for other DNA binding proteins and has been proposed to allow for bypass of obstacles along DNA such as other DNA-binding proteins and single nucleosomes during 1D damage search (24, 25, 43, 44). Compared to other proteins, the rate of diffusion of APE1 was extremely fast, likely facilitating efficient catalysis by this enzyme.

We found the ability of APE1 to search DNA in a 1D manner was dependent upon both the APE1 N-terminus and residue R177. While R177 provides a direct interaction with AP DNA by intercalating DNA strands at the AP-site (14, 15), a similar interaction with non-damaged DNA has not been previously observed, but our data points to this residue also being involved in retaining APE1 on non-damaged DNA during 1D scanning. The APE1 N-terminus was known to bind to specific non-damaged DNA sequences in the context of response elements (41, 42), but here we show that the APE1 N-terminus is necessary for binding to non-damaged DNA non-specifically to facilitate 1D damage search. Binding of the N-terminus to DNA during damage search may share properties with its binding to response elements, which is regulated by the acetylation of lysine residues within the APE1 N-terminus (42), and prompts interest in the acetylation states of APE1 during damage repair. However, we expect that APE1 binding to response element loci must be sequence specific and largely stationary, which we would not expect of APE1 binding during 1D diffusion along non-damaged DNA to search for damage (45, 46). While not definitive evidence, cryo-EM structures of APE1 bound to AP DNA have been unable to resolve the N-terminus (36), suggesting the N-terminus may have only transient interactions with non-damaged DNA. An intriguing hypothesis is that the unstructured nature of the N-terminus provides non-specific interactions with DNA that facilitate retention at the DNA when APE1 is undergoing micro disassociation events during 1D search mechanisms. While work remains to determine how R177 and the APE1 N-terminus specifically contribute to APE1 1D diffusion, our work demonstrates at the molecular level that these two features of APE1 contribute to search of non-damaged DNA via 1D diffusion.

APE1 recognized AP-sites in our study using both 1D and 3D diffusion, which is in agreement with a proposed theoretical model to improve the efficiency of repair protein search (26, 47). 1D diffusion allows for quick search of damage nearby an original non-specific location of DNA binding but takes longer to search sites far away from that original non-specific binding location (47). In comparison, 3D diffusion can quickly search different locations farther apart on the DNA in a relatively stochastic manner, but it may take longer to reach the damage site through random binding events (47). In this way, a balance between 1D and 3D diffusion is proposed to allow for efficient search of damage sites both far and near an original non-specific binding location (26, 47). Our results of APE1 searching for AP-sites seem to largely mirror this proposed model of 1D and 3D combinatory damage search, though APE1 did seem to utilize 3D search more often. This is consistent with the idea that 1D search is most effective over short distances (47). One in silico study estimated the length of DNA over which 1D search could be efficient to be a few hundred base pairs long (47), which is far shorter than the length of our 12.6 kbp DNA substrate. However, it is possible that some events we have categorized as 3D may demonstrate lengths of 1D search below the limit of the spatial resolution of our detector (< ∼300 bp). Similarly, we expect it is possible that APE1 could sample non-damaged DNA via 3D diffusion below the temporal resolution limit of our detector (< 5 ms). While these factors may limit the exact ratio of 1D vs 3D diffusion to the AP-site that we observe in this study, our findings are largely in line with the theoretical model that repair proteins like APE1 utilize a mix of both 1D and 3D diffusion to search DNA for damage.

Finally, regardless of the means of recognition, our results indicate that APE1 is retained at AP-sites upon their recognition. This would presumably facilitate repair, but retention can also potentially protect the toxic repair intermediate until the arrival of the next repair enzyme in the BER pathway (23, 48, 49). Interestingly, inhibition of repair by mutation of two key active site residues, D210 and E96, greatly increased the time APE1 was retained at AP-sites. However, the effects of mutation are not unique to simply preventing APE1 cleavage since all our experiments utilized a non-hydrolysable AP-site analog. Previous work has found mutation of residue E96 destabilizes the APE1 Mg^2+^ cofactor required for catalysis (14, 50), and this work along with our findings indicates Mg^2+^ likely regulates the dissociation of APE1 and AP DNA. Of note, this D210N/E96Q APE1 construct utilized in our study may provide a novel tool assessing the recruitment and localization of APE1 in previously difficult cellular imaging studies (37).

From the sum of our findings, we propose a novel model of APE1 search for AP-sites across the genome (Fig. 7). In this model, APE1 searches non-damaged DNA using both 1D and 3D diffusive mechanisms. 1D diffusion along non-damaged DNA is dependent upon the APE1 N-terminus and to a lesser extent the APE1 residue R177. Following APE1 recognition of AP-sites, APE1 is retained at the AP-site by residue R177 until the AP-site is appropriately cleaved. This cleavage is mediated by the residues D210 and E96 as well as the Mg^2+^ cofactor stabilized by residue E96. These residues and cofactor additionally regulate product release of cleaved DNA, allowing APE1 to dissociate from AP-sites after their cleavage. Overall, this work and the model proposed from it detail the process of APE1 damage search for the first time and lay a groundwork for investigating the damage search processes of other DNA repair enzymes at a molecular, mechanistic level.

**Figure 7:**
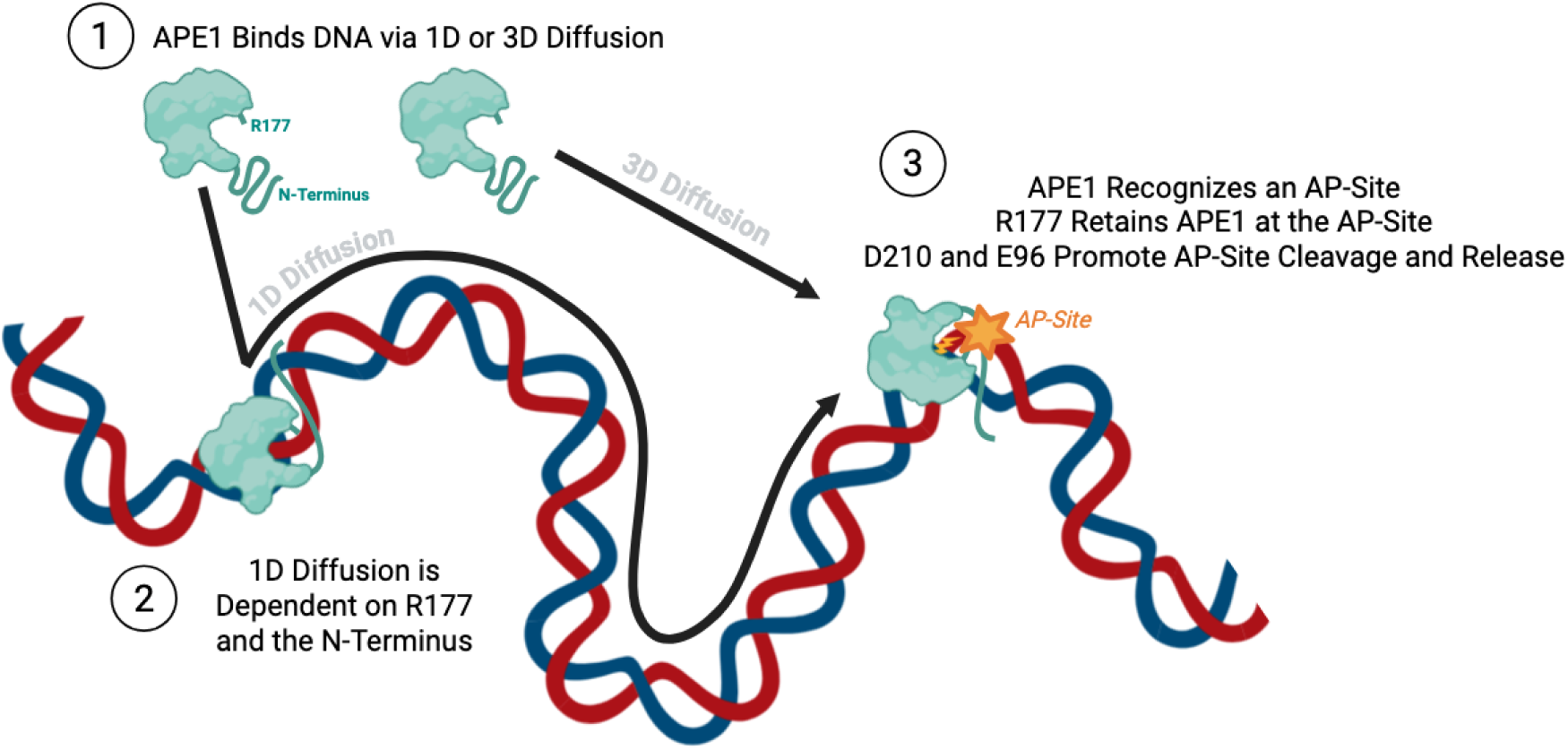
Proposed Model of APE1 Search for AP-Sites. (A) We propose that APE1 searches for AP-sites using both 1D and 3D diffusion. During 1D diffusive search, movement along non-damaged DNA is dependent upon the APE1 N-terminus, and to a lesser extent the residue R177. After AP-site recognition, APE1 is retained at the AP-site by residue R177 until cleavage. Cleavage is promoted by the residues D210 and E96, which promote release of APE1 from the AP-site following cleavage. This figure was created with BioRender.com.

## Materials and Methods

### APE1-Halo Protein Expression, Purification, and Fluorescent Labeling

*H. sapiens* APE1 and APE1 mutants with a C-terminal Halo-tag were expressed in NEB PLysY *E. coli* using a pFC30K vector. Cells were grown in 2XYT media at 37 °C and 200 rpm until reaching an OD of 0.6. Protein expression was induced following the initial growth phase and cells were grown for an additional period of 16 hrs at 20 °C. Cells were harvested and then lysed using a Fischer Scientific FB120 sonicator for 12 rounds of sonication lasting 30 seconds each at 90% power. Proteins were purified as previously described (36, 51). Briefly, lysate was filtered through a 0.45 um polyethersuflone membrane filter and then filtered lysate was purified using an AKTA-Pure FPLC with a Cytiva 5 mL Heparin HP affinity column. Fractions off the Heparin column containing APE1-Halo were pooled and purified with a Cytiva 5 mL Resource S cation exchange column on the ATKA. Finally, fractions off the Resource S column containing APE1-Halo were pooled, concentrated to 0.5 mLs and loaded onto a Cytiva HiPrep™ 16/60 Sephacryll71 S-200 HR gel filtration column on the AKTA. Purified protein was concentrated to 100 uM and stored at -80 °C.

APE1-Halo proteins were labeled with Janelia Fluorl71 Halo-Tag Dyes by combining 10 nmol of APE1-Halo with 20 nmols of a Janelia Fluorl71 H552 Halo-Tag Dye (22). Labeling reactions were incubated at room temperature for 1 hour, then labeled protein was purified using an ATKA-Pure FPLC with a Cytiva Superdexl71 200 Increase 10/30 GL gel filtration column. The efficiency of the labeling reactions was calculated by comparing the concentration of dye in the purified sample to the concentration of protein. Concentrations were calculated by measuring the absorbance of protein or dye using UV-vis spectroscopy and dividing the measured absorbance by the protein/dye’s extinction coefficient. Labeling efficiency for all APE1-Halo proteins in this study was > 90%. Purified fluorescently labeled proteins were concentrated to 10 μM and stored in dark conditions at -80 °C.

### DNA Tethering Substrates and Ligations

DNA substrates for CTFM experiments were prepared by ligating annealed DNA oligonucleotides purchased and purified by Integrated DNA Technologies (IDT) into DNA arms purchased and prepared by LUMICKS. LUMICKS’s two biotinylated DNA arms are 6298 base pairs of double stranded DNA each, and contain either an Atto647N fluorescent label and a GTTG single-strand overhang or an Atto488 fluorescent label and a TGGT single-strand overhang, respectively. Oligos purchased from IDT contained 54 nucleotides that when annealed incorporate a central 50 bp region of DNA and 4 bp overhangs complementing the LUMICKS arms single-strand overhangs. Oligonucleotides were annealed as previously described by resuspending lyophilized oligonucleotides in water to a final concentration of 10 μM, heating to 95 °C, and cooling at a rate of 1 °C per minute until reaching 4 °C (36, 51). Annealed oligonucleotides were ligated to LUMICKS DNA arms according to the manufacturer’s protocol with modifications. Briefly, 0.05 pmol of annealed oligonucleotides were incubated in 1x DNA Ligase Buffer with prepared arms and ligase as recommended by the manufacturer. Reactions were incubated at room temperature under dark conditions for 16 hours, then stored at 4C in siliconized microcentrifuge tubes for up to one month.

The 54-bp oligonucleotide sequences used in the ligation reactions were: 5’-CAACCGAGTGTCGCTGCCAACTAGGACTATCAAATGCTAGTGTTACAGGACTGC-3’ for the nondamaged template or 5’-CAACCGAGTGTCGCTGCCAACTAGGACXATCAAATGCTAGTGTTACAGGACTGC-3’ for a template where X denotes a non hydrolyzable tetrahydrofuran (THF) abasic site (52). The complement strand sequence was: 5’-ACCAGCAGTCCTGTAACACTAGCATTTGATAGTCCTAGTTGGCAGCGACACTCG-3’.

### Correlated Optical Tweezers with Fluorescence Microscopy (CTFM)

CTFM experiments were conducted using a LUMICKS C-Trapl71 Dymo equipped with two optical traps, a three-color (488, 561, 638 nm) confocal microscope, and the u-Flux™ microfluidics system. Prior to each experiment, the flow cell chamber was passivated with 0.1% BSA and a pluronic solution according to manufacturer instructions. Briefly, 0.5 mLs of 0.1% BSA solution was allowed to flow into the flow cell chamber at 0.3 bar for 30 mins, then 0.5 mLs of a 1:10 dilution of pluronic solution was allowed to flow into the flow cell chamber at 0.3 bar for 30 mins. Following passivation, a 1:1250 dilution to 1.5-1.9 µm streptavidin coated polystyrene beads was loaded into channel 1, a 1:250 dilution of DNA prepared as described above was loaded into channel 2, 250 pM fluorescently labeled APE1-Halo (or Δ1-42, R177A, D210N/E96Q APE1-Halo mutants) was loaded into channel 4, and buffer (20 mM HEPES, pH 7.5, 100 mM NaCl, 1 mM EDTA, 1 mM TCEP, and 0.1 mg/ml BSA, 0.45 µm FES-filtered) supplemented with 1 mM Trolox was loaded into channels 3 and 5. DNA and protein dilutions were prepared in the above buffer supplemented with Trolox, and beads were diluted in EMD Millipore sterile, WFI-quality, OmniPure water. Traps 1 and 2 were calibrated using 1.5-1.9 µm polystyrene bead templates and calibration was accepted if traps fell within 10% of average trap stiffness values of 0.5 pN at 20% overall trapping laser power. Following calibration, 1.5-1.9 µm streptavidin coated polystyrene beads were captured by placing traps 1 and 2 in the laminar flow path of channel 1. The trapped beads were then moved to the laminar flow path of channel 2 to catch the prepared biotinylated DNA substrate. Capture of a single DNA molecule was confirmed by fluorescent imaging and by force/distance measurements compared to an ideal force/distance curve simulated using an extensible worm-like chain (eWLC) model for a double strand DNA of 12.65 kbp (53, 54). Following successful DNA capture, DNA was stretched to 10 pN and exposed to fluorescently labeled APE1-Halo in the absence of flow. Kymographs from confocal imaging of APE1-Halo interactions with DNA were taken with a pixel size of 0.1, a scan line time of 10 ms, and all fluorescent laser sources at 5% power. Experiments were conducted at room temperature, averaging 25 °C according to microscope temperature sensors. Kymographs were obtained from a minimum of 3 unique dates corresponding to separate protein aliquots and 13 unique kymographs corresponding to unique strands of DNA until reaching the minimum number of binding events necessary to properly fit data (23).

### Kymograph Analysis and Particle Tracking

Kymographs were taken and exported as .h5 files using LUMICKS Bluelake software. Kymograph analysis was performed using Python v3.10.9 and the LUMICKS PyLake Python package v1.1.1, along with Anaconda v23.3.1 for environment management and Jupyter Notebook v6.5.4 as the IDE. Single-particle tracking was performed using the Kymotracking widget feature of PyLake, which utilizes the LUMICKS Greedy algorithm to determine tracking line centers. Customizable algorithm features included a search range of 0.35, a max length of 4, a max gap of 8 and a spot size of 0.80. We define “motile” particles as those where tracking of APE1-Halo particles displayed movement greater than 0.4 um within a period of 0.5 seconds.

### Diffusivity and Mean Squared Displacement Analysis

Following tracking, mean squared displacements (MSDs) and dwell times were calculated for each track. The MSD for each track was calculated using the equation:

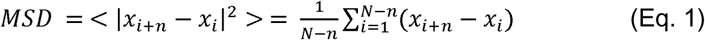

where x is the position at time frame i, n is the lag time of each consecutive frame, and N is the total number of points corresponding to the total number of frames in a track (24, 25, 28). The diffusion coefficient, D, for each track was determined by plotting MSDs over time and calculating the slope of the linear portion of the plot according to the following equation (24, 25, 28):

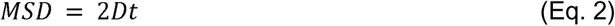

Each track was inspected for linearity and only tracks demonstrating an R^2^ > 0.8 were considered. Theoretical maximum limits of *D* for 1D diffusion were calculated for APE1-Halo on DNA for both sliding (aka rotational 1D diffusion) and hopping (aka non-rotational 1D diffusion) using Eq. 3, a form of the Stokes-Einstein-Sutherland equation (25, 28).

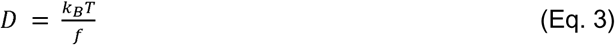

For Eq. 3, kB corresponds to the Boltzman constant commonly provided as 1.380649 x 10^-23^ joules/K, T is the temperature of the system in K, and f is defined as follows. When calculating the maximum theoretical limit of D of sliding 1D diffusion, f is defined as:

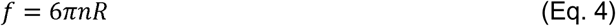

When calculating the maximum theoretical limit of D of sliding 1D diffusion, f is defined as:

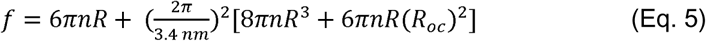

For Eqs. 4 and 5, n is the viscosity of a protein sliding along DNA in aqueous medium estimated as 9 x 10 ^ -10 pN s/nm^2^ (or 9 x 10 - 31 J s/nm^3^ in units adjusted to match the Boltzman constant) (55). For Eq. 5, 3.4 nm is the average length of 10-11 base pairs of DNA corresponding to one helical turn of DNA. R is the hydrodynamic radius of APE1-Halo, which was estimated at 4.41 nm using WinHYDROPRO (56). The structural model of APE1-Halo used for WinHYDROPRO calculations was generated using Alphafold (v 3.0) (57) and structures of individual APE1 and Halo molecules found in the PDB well (PDB IDs: 5DFF, 5UY1). R_oc_ is the distance of the hydrodynamic radius plus the distance of the protein to the center of the DNA helix. R_oc_ for APE1-Halo was estimated as 5.41 nm.

The average scan length was determined using the root mean squared displacement defined in the following equation:

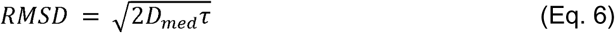

where *D_med_* is the median *D* determined as above, and τ corresponds to the binding lifetime determined as below (28, 58). The standard error the median reported as the standard error of the mean times √(π/2).

### Dwell Time and Gap Time Analysis

Dwell times were calculated for the length of each trace corresponding to a single binding event and were defined as:

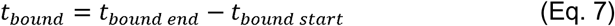

The binding lifetime (τ_bound_) was determined by Poisson regression defined as:

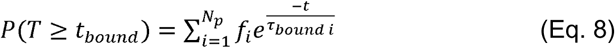

where P(T>t) is the probability that a dwell time, t_bound_, will be less than or equal to time T (23). In Eq 3.10., *N_p_* is total population of dwell times contributing to Poisson processes and *f_i_* is the fraction of events in each population of the Poisson regression. This probability is defined elsewhere as the cumulative residence time distribution (CRTD) (24) or 1 minus the cumulative distribution function (1-CDF) (25). The probability (provided here as 1-CDF) was plotted vs. time and fitted to a two-phase exponential decay curve, given by Eq. 9, to differentiate between populations of dwell times. The choice of a two-phase model was determined by F-tests comparing the fits of one, two, and three-phase models.

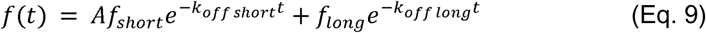

Binding lifetimes are related to off rate constants as defined by the equation *k_off_* = 1/τ_bound_ (23). Gap times were calculated as the time between consecutive binding events and were defined as (24, 25):

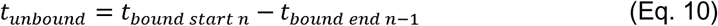

The gap interval also known as the inter-event interval, τ_free_, was determined by from all individual gap times by Poisson regression according to Eq. 11.

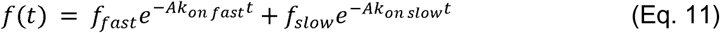

Gap intervals (τ_free_) are related to the on rate constants and enzyme concentration (A) as *k_on_* = 1/(A*τ_free_).

To calculate the binding affinity, we estimated a single on and single off rate constant for each APE1 variant at the AP-site that represented both the short and long populations referenced in Eqs. 9 & 11. An average rate constant calculated by weighing individual rate constants by the fraction comprising each population is described in Eq. 12. In Eq. 12, χ_short_ is the fraction given as a decimal of the population of events that are short-lived and χ_long_ is the fraction given as a decimal of the population of events that are long-lived.

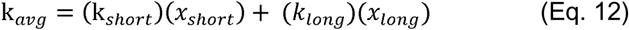

Binding affinity (*K*_D_) from single molecule measurements was then determined by Eq 3.13,

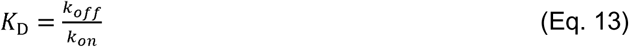

where off rate and on rate constants were determined from the binding lifetimes and gap intervals calculated to include all events measured under a given conditions.

### Electrophoretic Mobility Shift Assays (EMSAs)

The affinity of APE1-Halo and APE1-Halo mutants for either non-damaged or AP DNA was verified using EMSAs as previously described (36, 51). All DNA substrates used for EMSA experiments consisted of 30 base pairs of double-stranded DNA and contained a 5’ fluorescein (FAM) label for visualization. AP DNA contained a central non-hydrolysable phosphorothioate (THF) AP-site analog. DNA substrates were prepared by annealing oligonucleotides prepared by Integrated DNA Technologies (IDT). Annealing was performed by mixing 12 µM of the FAM- labeled template oligonucleotide with 10 µM of the unlabeled oligonucleotide complement in water and heating to 95 °C, then cooling at a rate of 1 °C per minute until reaching 4 °C. The oligonucleotide sequences used for non-damaged DNA were: 5’- *CGTTCGCTGATGCGCTCGACGGATCCGCAT-3’ and 5’-ATGCGGATCCGTCGAGCGCATCAGCGAACG-3’ (where * denotes the FAM-label). The oligonucleotide sequences used for AP DNA were: 5’-*CGTTCGCTGATGCGCXCGACGGATCCGCAT-3’ and 5’-ATGCGGATCCGTCGAGCGCATCAGCGAACG-3’ (where X denotes the THF AP-site analog) (52).

To conduct the EMSA experiments, various concentrations of APE1-Halo or APE1-Halo mutants were mixed with 2 nM of either non-damaged or AP-DNA and allowed to bind under non-hydrolysable conditions. For AP-DNA, concentrations of protein ranged from 0 to 20 nM for APE1-Halo, 0 to 250 nM for APE1^R177A^-Halo, 0 to 20 nM for APE1^Dead^-Halo, and 0 to 50 nM for APE1^Δ1-42^-Halo. For non-damaged DNA, concentrations of protein ranged from 0 to 500 nM for APE1-Halo, 0 to 5000 nM for APE1^R177A^-Halo, 0 to 625 nM for APE1^Dead^-Halo, and 0 to 5000 nM for APE1^Δ1-42^-Halo. The buffer conditions were as follows: 50 mM Tris, pH 8.0, 1 mM EDTA, 0.2 mg/mL BSA, 50 mg/mL sucrose, 0.5 mg/mL bromophenol blue, and 1 mM DTT. Binding reactions were incubated for 30 minutes to allow binding to approach equilibrium. Following incubation, samples were loaded onto a 59:1 polyacrylamide native gel. Gels were visualized using an Amersham Typhoon Imager. Bound and unbound fractions of DNA were quantified by densitometry using ImageJ. The fraction of bound DNA determined by ligand depletion was plotted against protein concentration, and the resulting curve was fit with Eq. 14 using Prism v9.1 to determine the apparent binding affinity (*K*_D App_).

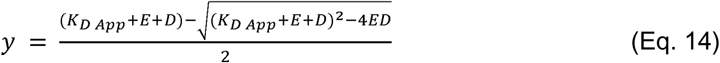

In Eq. 14, E is the enzyme concentration and D is the DNA concentration.

## Acknowledgments

We would like to thank Dr. Luke Lavis and his laboratory at the HHMI Janelia Research Campus for their provision of the Halo-tag ligands used in this study. We would also like to thank Mr. Anthony Vaughan for his assistance writing code to assist in calculating gap times. Finally, we thank previous and current members of LUMICKS including Matt Dilsaver, Nastaran Hadizadeh, Victor Su, and Edwin de Feijter. The authors utilized the LLM, ChatGPT, to edit portions of this manuscript for grammar and clarity. No LLMs were used for generation of any original content in this manuscript.

## Author Contributions

Kaitlin M. DeHart was responsible for project conceptualization, methodology, investigation, data curation, data analysis, data presentation, original manuscript drafting, and manuscript editing. Peyton N. Oden was responsible for investigation, data analysis, and manuscript review and editing. Tyler M. Weaver was responsible for project conceptualization, methodology, manuscript review and editing. Matt A. Schaich was responsible for methodology, data analysis, and manuscript review and editing. Ben Van Houtten was responsible for project supervision and manuscript review and editing. Bret D. Freudenthal was responsible for project supervision, project conceptualization, manuscript review and editing.

## Classification

Biological Sciences – Biophysics and Computational Biology

