## Supplemental Information for "A Dynamic Search Mechanism Enables APE1 to Identify AP-Sites in DNA"

**This PDF file includes:**

Supplemental Figure 1
Supplemental Tables 1-3

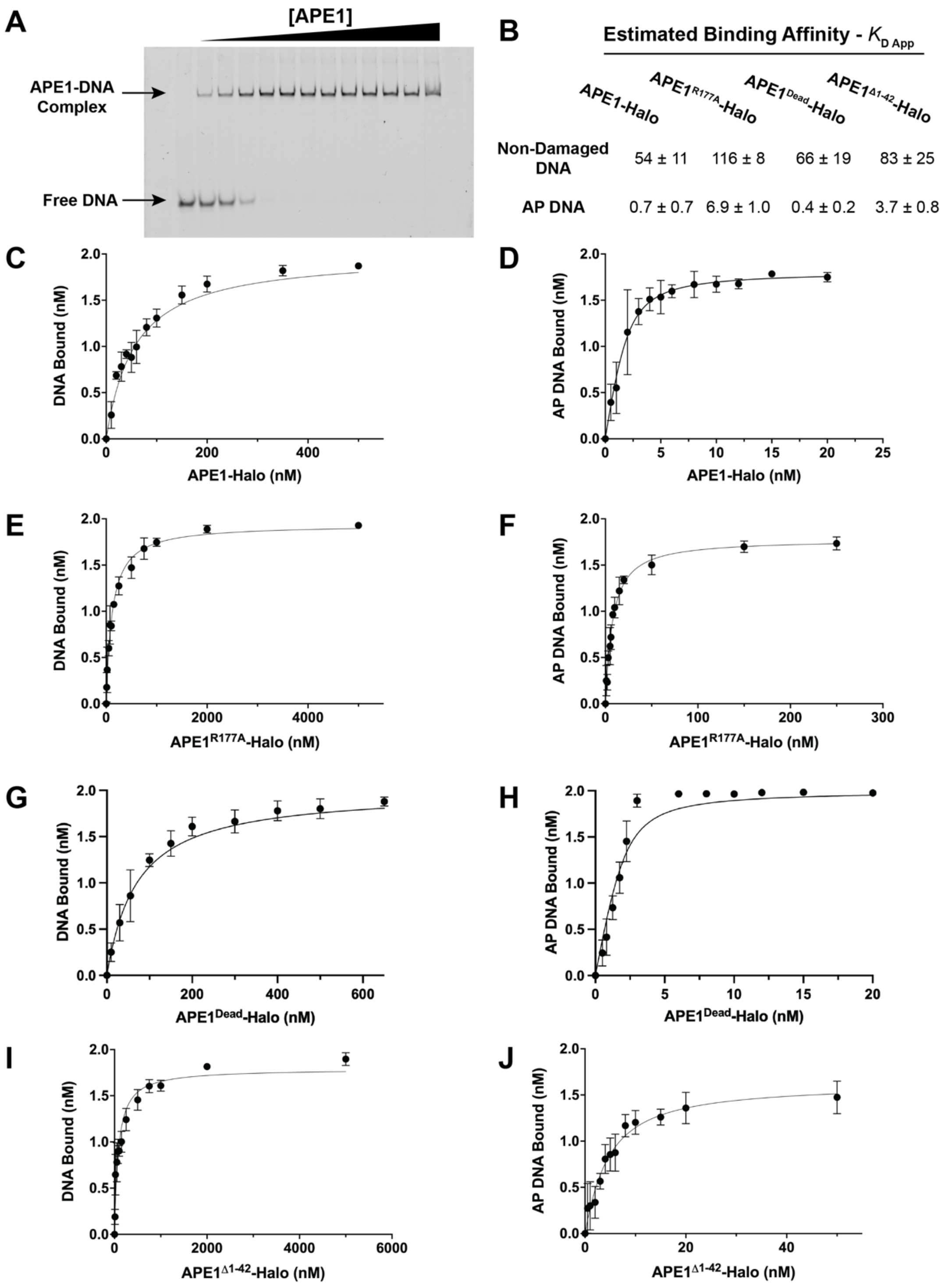


**Figure S1**

*Apparent Affinities of APE1-Halo and APE1-Halo Mutants Determined by EMSA*

*(A) Example of an EMSA gel demonstrating DNA bound by increasing an increasing concentration of APE1 enzyme. (B) Estimated apparent binding affinities ($K_{D\ App}$) of APE1-Halo and APE1-Halo mutants for both non-damaged and AP DNA as determined from EMSA experiments. Fits of the curves to determine the apparent binding affinities are shown for the following combinations of enzymes and DNA substrates: (C) APE1-Halo binding to non-damaged DNA. (D) APE1-Halo binding to AP DNA. (E) $APE1^{R177A}$-Halo binding to non-damaged DNA. (F) $APE1^{R177A}$-Halo binding to AP DNA. (G) $APE1^{Dead}$-Halo binding to non-damaged DNA. (H) $APE1^{Dead}$-Halo binding to AP DNA. (I) $APE1^{\Delta 1\text{-}42}$-Halo binding to non-damaged DNA. (J) $APE1^{\Delta 1\text{-}42}$-Halo binding to AP DNA.*

| | APE1-Halo | APE1$^{R177A}$-Halo | APE1$^{Dead}$-Halo | APE1$^{\Delta 1\text{-}42}$-Halo |
|---|---|---|---|---|
| **Motile Binding Events on Non-Damaged DNA** | | | | |
| Off Rate Constants | | | | |
| $k_{off\ short}$ (s$^{-1}$) | 3.2 ± 0.1 | | 2.1 ± 0.2 | |
| $k_{off\ long}$ (s$^{-1}$) | 0.38 ± 0.02 | | 0.84 ± 0.96 | |
| N | 315 | | 334 | |
| Distribution$_{bound}$ | 55.4% Short; 45.6% Long | | 88.5% Short; 11.5% Long | |
| **Stationary Binding Events at the AP-Site** | | | | |
| Off Rate Constants | | | | |
| $k_{off\ short}$ (s$^{-1}$) | 1.3 ± 0.1 | 10 ± 1 | 2.3 ± 0.1 | 1.4 ± 0.1 |
| $k_{off\ long}$ (s$^{-1}$) | 0.18 ± 0.01 | 1.1 ± 0.1 | 0.045 ± 0.001 | 0.62 ± 0.23 |
| N | 381 | 406 | 266 | 1158 |
| Distribution$_{bound}$ | 17.6% Short; 82.4% Long | 80.4% Short; 19.6% Long | 26.9% Short; 73.1% Long | 47.0% Short, 53% Long |
| On Rate Constants | | | | |
| $k_{on\ short}$ (nM$^{-1}$ s$^{-1}$) | 1.3 ± 0.1 | 1.0 ± 0.1 | 0.46 ± 0.02 | 2.0 ± 0.1 |
| $k_{on\ long}$ (nM$^{-1}$ s$^{-1}$) | 0.17 ± 0.01 | 0.28 ± 0.01 | 0.056 ± 0.005 | 0.85 ± 0.12 |
| N | 328 | 366 | 109 | 1086 |
| Distribution$_{free}$ | 45.2% Short; 54.8% Long | 60.1% Short; 39.9% Long | 57.9% Short; 42.1% Long | 83.0% Short; 17.0% Long |

**Table S1**

*Rate Constants Corresponding to Binding Lifetimes and Gap Intervals*

*Summary of off rate constants ($k_{off}$) and on rate constants ($k_{on}$), corresponding to binding lifetimes and gap intervals respectively, of APE1-Halo and APE1-Halo mutants. Rates are calculated for motile binding events on non-damaged DNA or for stationary binding events and the stationary portion of events exhibiting a mix of motile and stationary behavior at the AP-site. Rates are separated into short or long populations, and the proportion of each population is given by distribution of the total population of binding events (N).*

| | APE1-Halo | APE1$^{R177A}$-Halo | APE1$^{Dead}$-Halo | APE1 $^{\Delta 1\text{-}42}$-Halo |
|---|---|---|---|---|
| **Determined by CTFM** | | | | |
| $k_{off}$ ($s^{-1}$) | 0.38 ± 0.02 | 8.3 ± 0.4 | 0.64 ± 0.02 | 0.99 ± 0.05 |
| $k_{on}$ ($nM^{-1}$ $s^{-1}$) | 0.69 ± 0.01 | 0.73 ± 0.04 | 0.29 ± 0.03 | 1.5 ± 0.3 |
| $K_{D}$ (nM) | 0.55 ± 0.04 | 11 ± 1 | 2.2 ± 0.2 | 0.65 ± 0.12 |
| Determined by EMSA | | | | |
| $K_{D\ App}$ (nM) | 0.7 ± 0.7 | 6.9 ± 1.0 | 0.4 ± 0.2 | 3.7 ± 0.8 |

**Table S2**

**On and Off Rate Constants at the AP-site and Resulting Affinity Estimates**

*Summary of off and on rate constants ($k_{off}$ and $k_{on}$, respectively) of APE1-Halo, APE1$^{R177A}$-Halo, APE1$^{Dead}$-Halo, and APE1 $^{\Delta 1\text{-}42}$-Halo bound to an AP-site. Rates are a weighted average based on both populations of events as described in Table 1 (refer to Materials and Methods sections for more details). Binding affinity ($K_D$) was determined from off and on rates determined by CTFM experiments and separately from EMSA experiments.*

| | APE1-Halo | APE1$^{R177A}$-Halo | APE1$^{Dead}$-Halo | APE1$^{\Delta 1\text{-}42}$-Halo |
|---|---|---|---|---|
| **Kymographs Imaged on Non-Damaged DNA** | | | | |
| Total Binding Events | 359 | *NA* | 392 | *NA* |
| Total Unique Kymographs | 22 | *NA* | 16 | *NA* |
| Total Unique Dates Imaged | 3 | *NA* | 7 | *NA* |
| **Kymographs Imaged on AP DNA** | | | | |
| Total Binding Events | 699 | 491 | 1931 | 1231 |
| Total Unique Kymographs | 49 | 30 | 152 | 20 |
| Total Unique Dates Imaged | 7 | 5 | 23 | 3 |

**Table S3**

*Table of Kymograph Replicates*

*Summary of imaging replicates for APE1-Halo and each APE1-Halo mutant on both non-damaged and AP DNA. The total number of traces were acquired from kymographs where each kymograph corresponded to a new strand of DNA imaged (analogous to biological replicates). Reagents were prepared from stocks for each conditions over the course of at least separate dates (analogous to technical replicates).*
